# Forsythoside B attenuates neuropathic pain by suppressing USP7/HIF-1α-mediated microglial glycolytic reprogramming

**DOI:** 10.64898/2026.09.20.752649

**Authors:** Zhen dai, Hao Wang, Yang Yanfeng

## Abstract

Neuropathic pain involves spinal microglial activation and metabolic changes. Forsythoside B (FB) has anti-inflammatory effects, but its role in neuropathic pain and microglial glycolysis remains unclear. We investigated the effects and mechanisms of FB in mice with chronic constriction injury (CCI) and in LPS-stimulated BV2 microglia. FB reduced mechanical allodynia and thermal hyperalgesia in CCI mice. FB also reduced the spinal levels of HIF-1α, HK2, PKM2, LDHA, and CD86. It decreased the proportion of IBA1+/CD86+ cells and the levels of IL-1β, IL-6, and TNF-α. In LPS-stimulated BV2 microglia, FB reduced glycolysis-related proteins, proinflammatory markers, and cytokine release. FB also reduced glycolytic activity and partly restored basal mitochondrial respiration. FB did not change Hif1a mRNA levels. However, it increased HIF-1α ubiquitination and promoted its proteasome-dependent degradation. HIF-1α overexpression partly reversed the effects of FB on glycolytic enzymes, CD86 expression, and proinflammatory cytokine production. Immunoprecipitation followed by mass spectrometry and UbiBrowser analysis identified USP7 as a candidate HIF-1α-associated deubiquitinase. Molecular docking and cellular thermal shift assays supported a possible interaction between FB and USP7. FB reduced the association between USP7 and HIF-1α. USP7 knockdown produced effects similar to those of FB on HIF-1α ubiquitination and downstream glycolytic and inflammatory responses. Overall, FB alleviated neuropathic pain and reduced proinflammatory microglial responses. These effects may involve the promotion of HIF-1α degradation through modulation of the USP7-HIF-1α regulatory axis.

**Graphical abstract:** 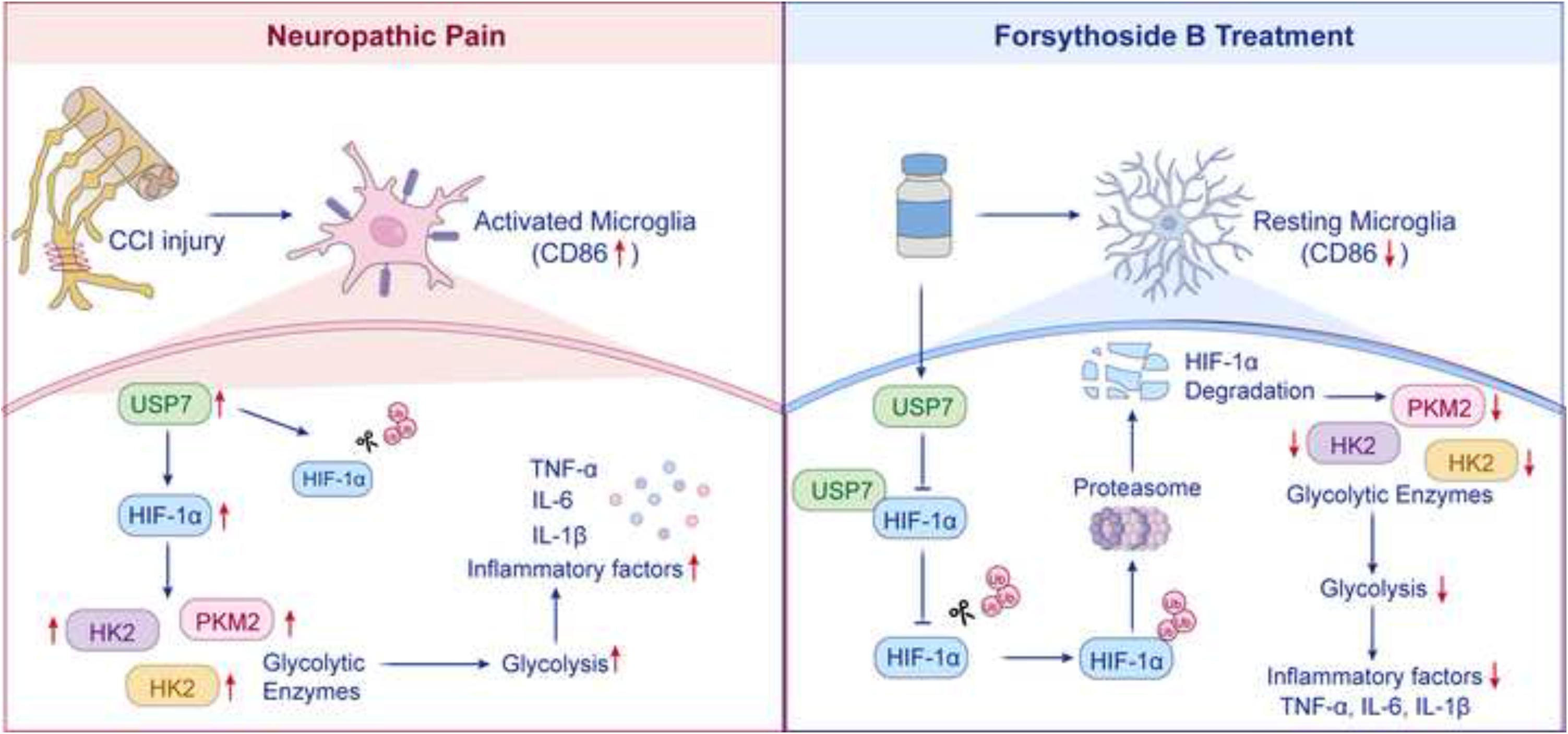

## 1. Introduction

Neuropathic pain (NP) is a chronic condition caused by a lesion or disease affecting the somatosensory nervous system. It can result from diabetes, infection, traumatic injury, or neurological disease [1,2]. NP affects an estimated 7–10% of the global population and places a substantial burden on patients, particularly older adults [3]. Persistent symptoms can impair daily activities and reduce quality of life. Patients with NP also commonly experience anxiety and depression. Severe NP has been associated with an increased risk of suicide [4]. Common clinical features include spontaneous pain, hyperalgesia, and mechanical allodynia [5].

Microglial activation in the spinal dorsal horn plays an important role in the development and persistence of neuropathic pain [6–8]. After nerve injury, microglia rapidly transition from a surveillant, ramified morphology to an activated amoeboid state. This change is accompanied by increased release of pro-inflammatory mediators , which can enhance nociceptive signaling [9]. Microglial activation is closely associated with metabolic remodeling. Resting microglia mainly depend on mitochondrial oxidative phosphorylation. Activated microglia, in contrast, rely more on aerobic glycolysis and show reduced tricarboxylic acid cycle activity [10]. Hypoxia-inducible factor 1α (HIF-1α) is an important transcriptional regulator in this process. It connects inflammatory signaling with glycolytic programming by inducing the expression of glycolysis-related genes [11]. Sustained glycolytic activity and metabolite accumulation may further promote a proinflammatory phenotype and impair phagocytosis. These changes can contribute to persistent neuroinflammation [12]. Therefore, targeting microglial metabolic reprogramming may provide a potential strategy for treating neuropathic pain.

Ubiquitin-specific protease 7 (USP7) is a deubiquitinase that controls the stability and activity of multiple proteins. Recent studies have linked USP7 to microglia-mediated neuroinflammation. USP7 inhibition promotes the ubiquitin-dependent degradation of Kelch-like ECH-associated protein 1 (Keap1), activates nuclear factor erythroid 2-related factor 2 (Nrf2) signaling, and reduces proinflammatory microglial activation [13]. Consistent with these findings, targeting the USP7/Keap1/Nrf2 pathway reduces spinal inflammation and pain hypersensitivity after peripheral nerve injury [14]. These results suggest that USP7 may contribute to neuropathic pain. USP7 may also regulate HIF-1α stability. It can interact with HIF-1α and remove its ubiquitin modifications, thereby limiting HIF-1α degradation and increasing its protein stability [15,16]. Stable HIF-1α can enhance glycolysis and promote proinflammatory microglial activation in experimental models of neuropathic pain [8]. These findings suggest that USP7 may control microglial immunometabolic remodeling by regulating HIF-1α stability.

Although pharmacotherapy remains the mainstay of neuropathic pain management, its clinical benefit is often limited by modest efficacy and treatment-related adverse effects [17]. First-line anticonvulsants and antidepressants may cause dizziness, somnolence, headache, and hypertension, which can compromise treatment adherence and long-term symptom control [7]. These limitations have prompted interest in plant-derived natural products with multi-target anti-inflammatory and analgesic activities. Forsythoside B (FB), a phenylethanoid glycoside isolated from *Forsythia suspensa*, has demonstrated antioxidant, anti-apoptotic, anti-inflammatory, and pro-regenerative effects in experimental models [18]. Previous studies have shown that FB reduces cognitive impairment and neuroinflammation in models of Alzheimer’s disease [19] and attenuates apoptosis and inflammation after spinal cord injury [20]. FB also alleviates inflammatory pain and suppresses microglial and astrocytic activation [21]. However, whether FB exerts analgesic effects in neuropathic pain and whether microglial metabolic reprogramming contributes to these effects remain unclear. Therefore, using a chronic constriction injury (CCI) model and LPS-stimulated BV2 microglia, we investigated the analgesic and anti-neuroinflammatory effects of FB and examined whether FB modulates microglial glycolytic reprogramming through the USP7/HIF-1α axis.

## 2. Materials and Methods

### 2.1. Animals and experimental design

Forsythoside B (FB; HY-N0029; purity ≥ 99.99%) was purchased from MedChemExpress (MCE, USA). Male C57BL/6J mice (6-8 weeks old; 18-22 g) were obtained from the Experimental Animal Center of Jinzhou Medical University. Mice were housed under controlled conditions (22 ± 1 °C; 12-h light/dark cycle) with free access to food and water. All animal procedures were approved by the Animal Ethics Committee of the First Affiliated Hospital of Jinzhou Medical University (Approval No. JZMULL2025240).

Mice were randomly assigned to five groups (n = 6 per group): Sham, CCI, CCI + FB-L (10 mg/kg), CCI + FB-M (20 mg/kg), and CCI + FB-H (40 mg/kg). FB was dissolved in saline and administered by intraperitoneal injection once daily from postoperative day 1 for 14 consecutive days. Mice in the Sham and CCI groups received an equivalent volume of saline.

### 2.2. Establishment of the CCI model

Mice were anesthetized with 1% sodium pentobarbital (50 mg/kg, i.p.). The right sciatic nerve was exposed at the mid-thigh level and loosely ligated with three 4-0 Prolene sutures spaced approximately 1 mm apart. Each ligature was tightened until a brief twitch of the ipsilateral hind limb was observed, indicating mild nerve constriction without complete occlusion. Sham-operated mice underwent identical anesthesia and sciatic nerve exposure without ligation.

### 2.3. Behavioral assessment

Mechanical and thermal nociceptive sensitivities were assessed before surgery (day 0) and on postoperative days 3, 7, 10, and 14 by an investigator blinded to group allocation. Mechanical sensitivity was evaluated using von Frey filaments as previously described [22]. After a 30-min habituation period, calibrated filaments were applied perpendicularly to the plantar surface of the right hind paw. The paw withdrawal threshold (PWT) was defined as the lowest force that evoked a withdrawal response in at least 5 of 10 applications.

Thermal sensitivity was assessed using a hot-plate assay based on a published protocol with minor modifications [23]. After 10 min of acclimation to the testing environment, mice were placed on a heated surface maintained at 52 ± 0.2 °C. Paw withdrawal latency (PWL) was recorded as the time to the first nocifensive response, including hind-paw licking, flicking, or jumping. A cutoff time of 20 s was applied to prevent tissue injury.

### 2.4. Measurement of inflammatory cytokines

Cytokine concentrations (IL-1β, IL-6, and TNF-α) in spinal cord homogenates and BV2 culture supernatants were determined with commercially available ELISA kits (Jianglai Biotech, Shanghai, China).

### 2.5. Cell culture and treatments

The murine BV2 microglial cell line was obtained from the Cell Bank of the Chinese Academy of Sciences. Cells were maintained in Dulbecco’s modified Eagle’s medium (DMEM; Solarbio, Beijing, China) supplemented with 10% fetal bovine serum and 1% penicillin-streptomycin at 37 °C in a humidified atmosphere containing 5% CO₂. For in vitro experiments, cells were assigned to the following groups: Control, LPS, LPS + FB (2.5 μM), and LPS + FB (5 μM). FB was added 2 h before LPS stimulation. Control and LPS groups received an equivalent volume of vehicle during the pretreatment period. Cells were then exposed to LPS (1 μg/mL; Beyotime, Shanghai, China) for 24 h, whereas control cells received culture medium alone. Culture supernatants and cell lysates were subsequently collected for downstream analyses.

### 2.6. Cell viability assay

Cell viability was assessed using a Cell Counting Kit-8 (CCK-8; Beyotime, Shanghai, China). BV2 cells were treated with the indicated concentrations of FB for 24 h. Subsequently, 10 μL of CCK-8 reagent was added to each well, followed by incubation at 37 °C for 2 h. Absorbance was measured at 450 nm using a microplate reader (BioTek, USA). Cell viability was expressed relative to that of the vehicle-treated control group.

### 2.7. Cell transfection

USP7-targeting siRNAs, negative-control siRNA, the HIF-1α-overexpression plasmid (pcDNA3.1-CMV-HIF-1α), and the corresponding empty vector were obtained from GenePharma (Shanghai, China). BV2 cells at 60-70% confluence were transfected using Lipofectamine 3000 (Invitrogen, USA) according to the manufacturer’s instructions. After 48 h, transfected cells were subjected to the indicated treatments and subsequent analyses. The sequences of the USP7-targeting siRNAs are provided in Supplementary Table 1.

### 2.8. Extracellular flux assay (Seahorse)

BV2 cells were seeded in Seahorse XF24 microplates at 5 × 10⁴ cells per well and subjected to the indicated treatments. ECAR and OCR were measured using a Seahorse XF24 Analyzer (Agilent Technologies, USA).

For ECAR analysis, cells were equilibrated in Seahorse XF base medium containing 2 mM glutamine for 1 h at 37 °C. Glucose, oligomycin, and 2-deoxy-D-glucose were injected sequentially to determine glycolysis, glycolytic capacity, and glycolytic reserve. For OCR analysis, cells were equilibrated in Seahorse XF assay medium containing 10 mM glucose, 2 mM glutamine, and 1 mM sodium pyruvate for 1 h at 37 °C. Oligomycin, FCCP, and rotenone/antimycin A were injected sequentially to determine mitochondrial respiratory parameters.

### 2.9. Western blot analysis

BV2 cells and L4-L6 spinal cord tissues were lysed in RIPA buffer (Epizyme, Shanghai, China) supplemented with protease inhibitors on ice for 30 min. Lysates were centrifuged to remove cellular debris, and protein concentrations were determined using a BCA protein assay kit (Epizyme). Equal amounts of protein were separated by SDS-PAGE and transferred onto methanol-activated polyvinylidene fluoride membranes using a wet-transfer system. Membranes were blocked with 5% non-fat milk in TBST for 2 h at room temperature and incubated overnight at 4 °C with primary antibodies against USP7 (1:1,000; Proteintech), HIF-1α (1:1,000; Abcam), HK2 (1:1,000; Proteintech), PKM2 (1:1,000; Proteintech), LDHA (1:1,000; Proteintech), CD86 (1:1,000; Cell Signaling Technology), and β-actin (1:10,000; Proteintech). After washing with TBST, membranes were incubated with HRP-conjugated secondary antibodies (1:10,000; Proteintech) for 1 h at room temperature. Protein signals were visualized using an enhanced chemiluminescence reagent and captured using a ChemiDoc imaging system (Bio-Rad).

### 2.10. Co-immunoprecipitation

BV2 cells were lysed in RIPA buffer, and the lysates were centrifuged to remove cellular debris. For each immunoprecipitation, 500 μg of total protein was incubated with 2 μg of control IgG or the indicated primary antibody, including anti-USP7 or anti-HIF-1α antibody, overnight at 4 °C with gentle rotation. Protein A/G agarose beads (40 μL; Beyotime, Shanghai, China) were then added and incubated for an additional 3 h at 4 °C. The beads were washed with lysis buffer, and bound proteins were eluted by boiling in 1× SDS loading buffer before Western blot analysis

### 2.11. Protein stability assay

HIF-1α protein stability was assessed using a cycloheximide (CHX) chase assay. Following the indicated treatments, BV2 cells were incubated with CHX (100 μg/mL; MedChemExpress, USA) to inhibit de novo protein synthesis. Cells were collected at 0, 3, 6, and 9 h after CHX treatment, and total protein lysates were prepared for Western blot analysis. HIF-1α protein levels at each time point were quantified by densitometry and normalized to the level at 0 h to assess protein degradation kinetics.

### 2.12. Assessment of the HIF-1α degradation pathway

To determine the pathway involved in FB-induced HIF-1α degradation, LPS-stimulated BV2 cells were treated with FB in the presence or absence of the proteasome inhibitor MG132 (10 μM; MedChemExpress, USA) or the lysosomal inhibitor chloroquine (CQ; 50 μM; MedChemExpress, USA). HIF-1α protein levels were subsequently assessed by Western blotting.

### 2.13. Immunoprecipitation-mass spectrometry analysis

To identify HIF-1α-interacting proteins, BV2 cells were stimulated with LPS (1μg/mL) for 24 h. Cell lysates were subjected to immunoprecipitation using an anti-HIF-1α antibody. The immunoprecipitated proteins were analyzed by liquid chromatography-tandem mass spectrometry at Majorbio Bio-Pharm Technology Co., Ltd. (Shanghai, China). Proteins identified by mass spectrometry were screened for E3 ubiquitin ligases and deubiquitinating enzymes and compared with HIF-1α regulators listed in UbiBrowser 2.0.

### 2.14. Molecular docking simulation

Molecular docking was performed to investigate the potential binding mode of FB with USP7. The crystal structure of USP7 (PDB ID: 5VSB) was obtained from the RCSB Protein Data Bank (https://www.rcsb.org/). The three dimensional structure of FB was retrieved from the PubChem database (https://pubchem.ncbi.nlm.nih.gov/). The protein and ligand were prepared using standard procedures before docking analysis with AutoDock Vina. The predicted binding mode was visualized using PyMOL and Discovery Studio Visualizer.

### 2.15. Cellular thermal shift assay (CETSA)

The cellular thermal shift assay was performed to assess the effect of FB on USP7 thermal stability. BV2 microglia were treated with vehicle or FB (5 μM) under the indicated experimental conditions. Cells were collected, resuspended in phosphate-buffered saline supplemented with protease inhibitors, and divided into equal aliquots. The aliquots were heated at 40, 45, 50, 55, 60, or 65 °C, followed by cooling on ice. Cell lysates were then subjected to freeze–thaw cycles and centrifuged to remove precipitated proteins. The soluble fractions were collected, and residual USP7 protein levels were determined by western blotting. USP7 signals were normalized to β-actin and expressed relative to the corresponding sample heated at 40 °C.

### 2.16. Immunofluorescence staining

Spinal cord sections and BV2 cells were fixed with 4% paraformaldehyde (Beyotime, Shanghai, China), permeabilized with 0.5% Triton X-100 (Beyotime), and blocked with 5% bovine serum albumin. Samples were incubated overnight at 4 °C with the following primary antibodies: mouse anti-IBA1 (1:200; Abcam), rabbit anti-USP7 (1:200; Sanying, Wuhan, China), rabbit anti-CD86 (1:200; Cell Signaling Technology), and rabbit anti-HIF-1α (1:200; Abcam). After washing, samples were incubated with the appropriate fluorophore-conjugated secondary antibodies (Epizyme, Shanghai, China) for 1 h at room temperature in the dark. Nuclei were counterstained with DAPI. Fluorescence images were acquired using an inverted fluorescence microscope.

### 2.17. Real-time quantitative PCR

Total RNA was extracted from L4-L6 spinal cord tissues and cultured BV2 cells using TRIzol reagent and subsequently purified using a commercial RNA purification kit (EZBioscience, China). Complementary DNA was synthesized by reverse transcription. Quantitative PCR was performed using SYBR Green Master Mix (Vazyme, China) on a real-time PCR system. The amplification protocol consisted of an initial denaturation step at 95 °C for 2 min, followed by 40 cycles of 95 °C for 10 s, 50 °C for 30 s, and 72 °C for 30 s. Relative mRNA expression was calculated using the 2^−ΔΔCt method. Primer sequences are provided in Table 1.

**Table 1.**
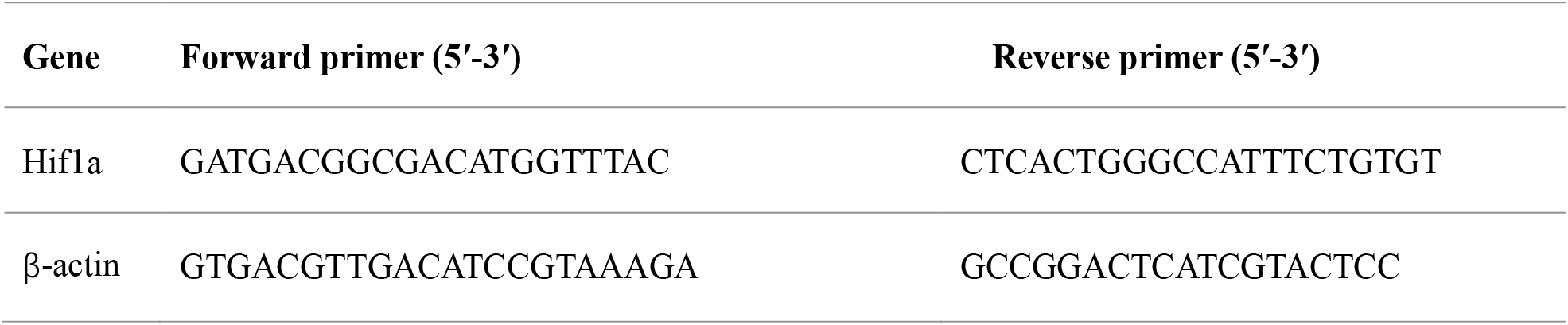
The list of primer sequences.

### 2.18. Statistical analysis

Data are presented as mean ± SEM. Two-group comparisons were performed using an unpaired two-tailed Student’s t-test. Comparisons among multiple groups were analyzed by one-way ANOVA followed by Tukey ’ s multiple comparisons test. Behavioral data measured repeatedly over time were analyzed by two-way repeated-measures ANOVA (group × time) followed by Sidak’s multiple comparisons test. Statistical analyses were conducted using GraphPad Prism 10, and statistical significance was defined as *P* < 0.05.

## 3. Results

### 3.1. FB alleviates pain hypersensitivity and spinal neuroinflammation in CCI mice

To assess the effects of FB in CCI mice, we measured mechanical and thermal nociceptive sensitivity. Compared with Sham mice, CCI mice showed lower PWT and PWL from postoperative day 3 onward, indicating mechanical allodynia and thermal hyperalgesia, respectively (Fig. 1A,B). Medium- and high-dose FB increased both PWT and PWL from postoperative day 7 to day 14 compared with CCI mice. Low-dose FB produced no consistent improvement in pain hypersensitivity.

**Fig. 1.**
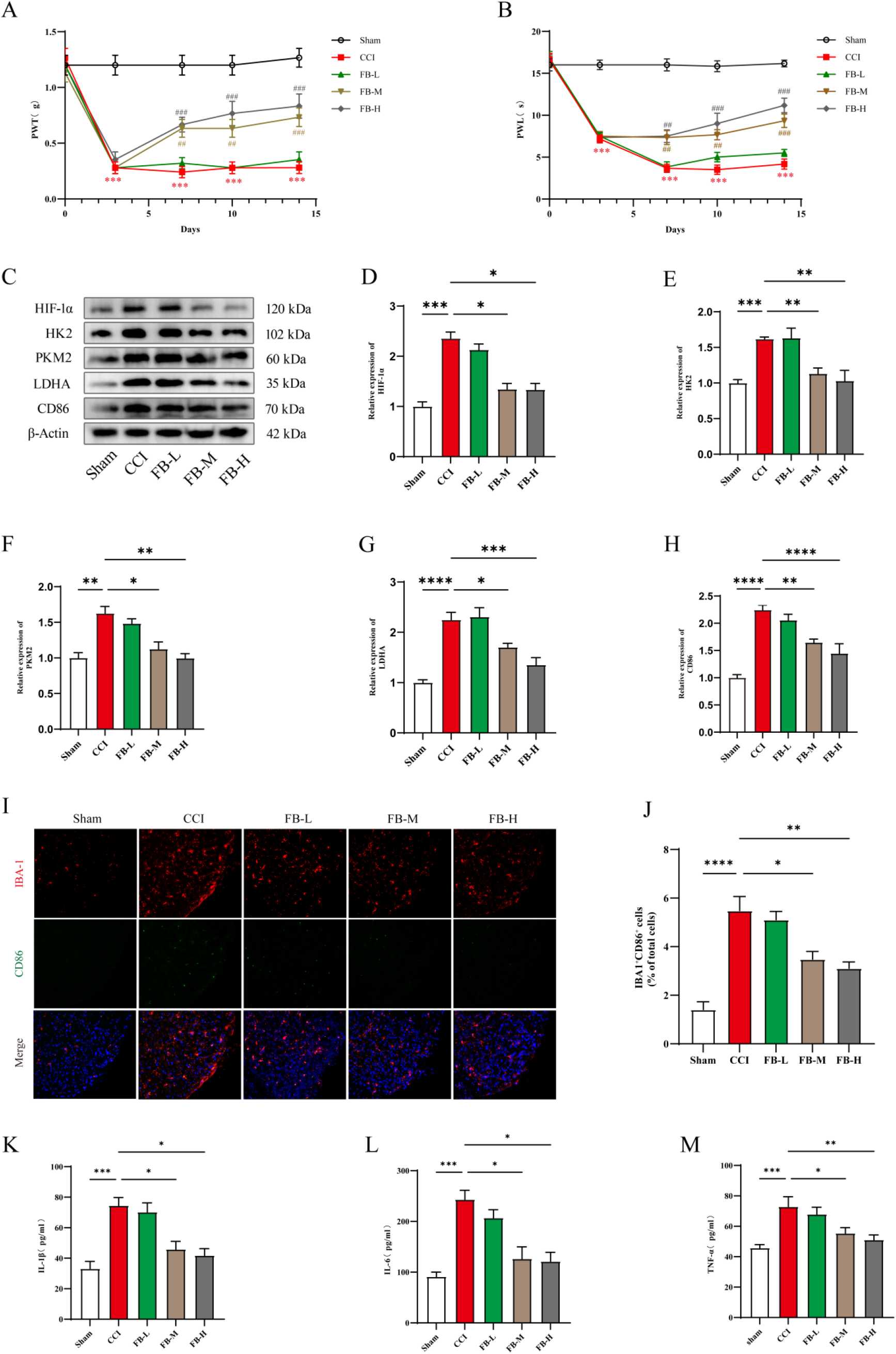
FB alleviates pain hypersensitivity, spinal neuroinflammation, and changes in proteins involved in glycolysis in CCI mice. (A) PWT at the indicated time points after surgery. (B) PWL at the indicated time points after surgery. (C) Representative western blot images of HIF-1α, HK2, PKM2, LDHA, CD86, and β-actin in L4-L6 spinal cord tissues. (D-H) Quantification of HIF-1α, HK2, PKM2, LDHA, and CD86 protein levels, respectively, normalized to β-actin. (I) Representative immunofluorescence images of IBA1 (red), CD86 (green), and DAPI (blue) in the spinal dorsal horn. Scale bar = 100 μm. (J) Quantification of IBA1+/CD86+ cells as a percentage of total cells. (K-M) IL-1β, IL-6, and TNF-α levels, respectively, in L4-L6 spinal cord tissues determined by ELISA. FB-L, FB-M, and FB-H represent low, medium, and high doses of FB, respectively. Data are presented as mean ± SEM (n = 6 mice per group). *P* < 0.05, *P* < 0.01, *P* < 0.001, *P* < 0.0001; #*P* < 0.05, ##*P* < 0.01, ###*P* < 0.001.

We then examined inflammatory and metabolic changes in the spinal cord. CCI increased the protein levels of HIF-1α, HK2, PKM2, LDHA, and CD86 in L4–L6 spinal cord tissues compared with Sham mice (Fig. 1C–H). Medium- and high-dose FB reduced these protein levels, whereas low-dose FB had limited effects. CCI also increased the proportion of IBA1+/CD86+ cells in the spinal dorsal horn. FB reduced this proportion at medium and high doses (Fig. 1I,J). ELISA further showed that CCI increased the levels of IL-1β, IL-6, and TNF-α in L4–L6 spinal cord tissues. Medium- and high-dose FB reduced these cytokine levels (Fig. 1K–M). Overall, FB reduced pain hypersensitivity and spinal inflammatory responses in CCI mice. These effects were accompanied by lower levels of HIF-1α and glycolysis-related proteins.

### 3.2. FB suppresses LPS-induced inflammatory activation and HIF-1α-associated glycolytic changes in BV2 microglia

To determine whether the effects of FB observed in vivo could be reproduced in microglia, we used LPS-stimulated BV2 cells as an in vitro model of microglial inflammation. CCK-8 assays showed that FB at concentrations of 2.5–40 μM did not significantly affect BV2 cell viability, whereas 80 μM FB reduced cell viability (Fig. 2A). Therefore, 2.5 and 5 μM FB were used in subsequent experiments. Western blotting showed that LPS increased the protein levels of HIF-1α, the glycolysis-related enzymes HK2, PKM2, and LDHA, and the proinflammatory marker CD86 compared with the control group (Fig. 2B–G). FB treatment reduced several of these LPS-induced changes, with generally stronger effects at 5 μM. Immunofluorescence analysis further showed that LPS increased CD86 expression, whereas 5 μM FB reduced this increase (Fig. 2H,I). ELISA showed that LPS also increased the levels of IL-1β, IL-6, and TNF-α in the culture supernatant. Treatment with 5 μM FB reduced the release of these proinflammatory cytokines (Fig. 2J–L). Overall, FB reduced LPS-induced inflammatory activation in BV2 microglia and decreased the levels of HIF-1α and glycolysis-related proteins.

**Fig. 2.**
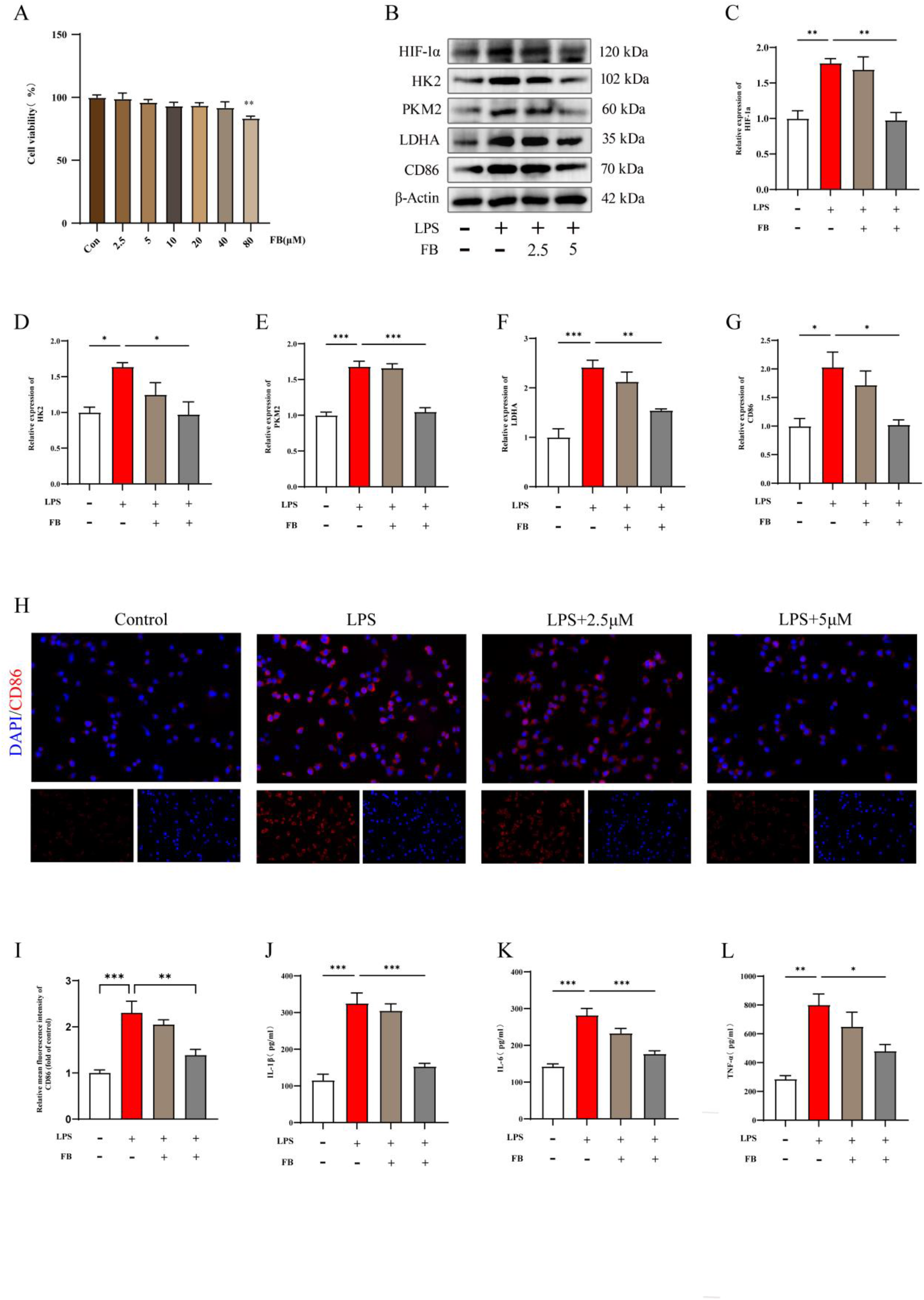
FB suppresses LPS-induced inflammatory responses and HIF-1α-associated glycolytic protein expression in BV2 microglia. (A) Cell viability of BV2 cells treated with the indicated concentrations of FB, as assessed by the CCK-8 assay. (B) Representative Western blot images of HIF-1α, HK2, PKM2, LDHA, and CD86 in LPS-stimulated BV2 cells treated with or without FB. (C–G) Quantitative analysis of HIF-1α, HK2, PKM2, LDHA, and CD86 protein levels, respectively, normalized to β-actin. (H) Representative immunofluorescence images showing CD86 (red) and DAPI (blue) staining in BV2 cells. Scale bar = 50 μm. (I) Quantification of relative mean CD86 fluorescence intensity. (J–L) IL-1β, IL-6, and TNF-α levels in the culture supernatants measured by ELISA. Data are presented as the mean ± SEM (n = 3). \**P* < 0.05, \*\**P* < 0.01, and \*\*\**P* < 0.001.

### 3.3. FB attenuates LPS-induced glycolytic activation and partially restores basal mitochondrial respiration in BV2 microglia

To characterize the effects of FB on cellular metabolism, we assessed glycolytic activity and mitochondrial respiration by measuring the extracellular acidification rate (ECAR) and oxygen consumption rate (OCR). LPS stimulation increased glycolysis, glycolytic capacity, and glycolytic reserve in BV2 cells (Fig. 3A–D). Treatment with 5 μM FB reduced these LPS-induced increases, although the glycolytic parameters remained higher than those in the control group. OCR analysis showed that LPS stimulation reduced basal respiration compared with the control group. FB treatment partially restored basal respiration, although the level remained lower than that in the control group (Fig. 3E,F). FB did not significantly improve maximal respiration, ATP production, or spare respiratory capacity compared with the LPS group (Fig. 3G–I).

**Fig. 3.**
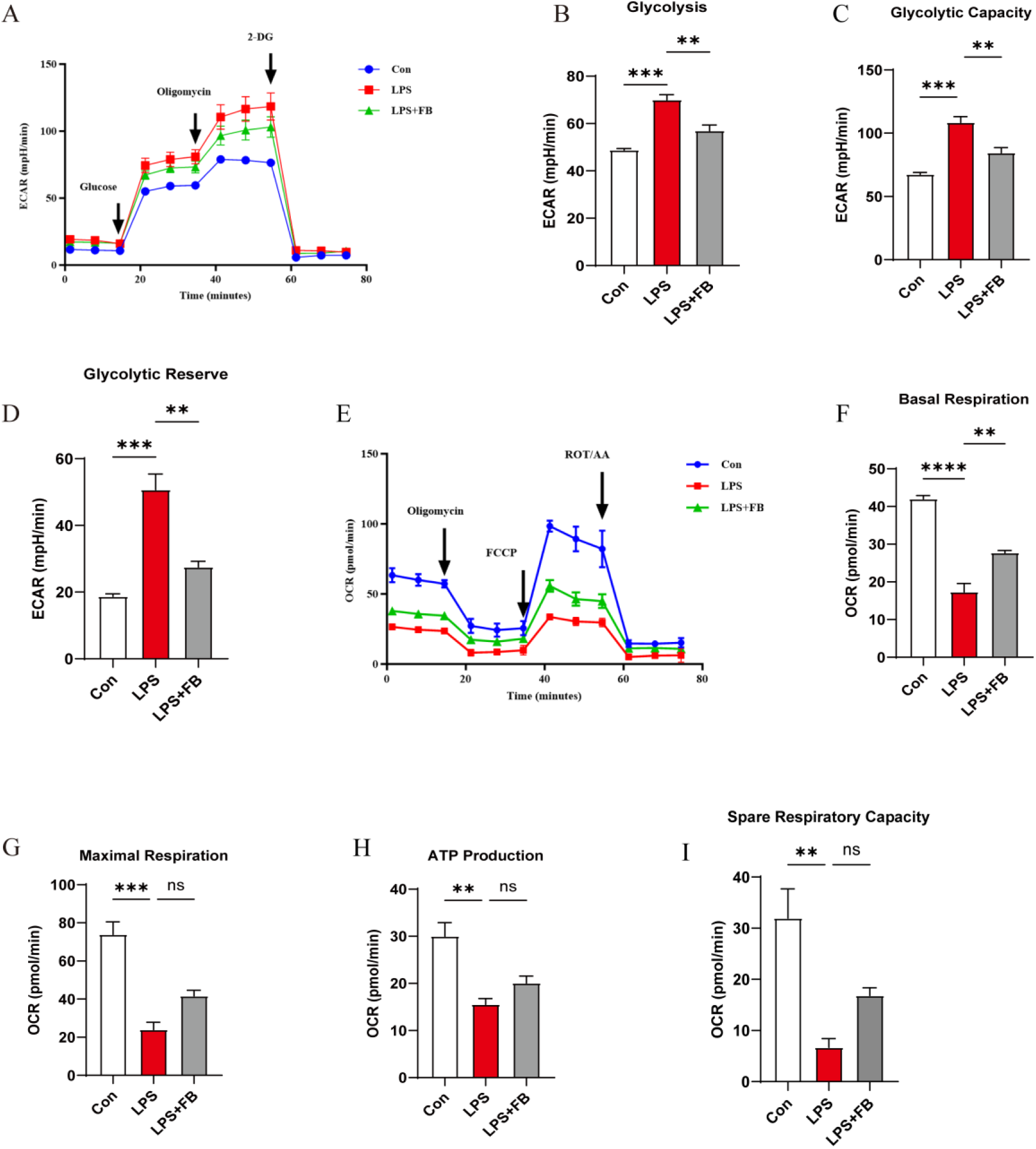
FB modulates glycolysis and mitochondrial respiration in LPS-stimulated BV2 microglia. (A) Representative real-time ECAR traces after sequential injection of glucose, oligomycin, and 2-deoxy-D-glucose (2-DG). (B–D) Quantification of glycolysis, glycolytic capacity, and glycolytic reserve, respectively. (E) Representative real-time OCR traces after sequential injection of oligomycin, FCCP, and rotenone/antimycin A. (F–I) Quantification of basal respiration, maximal respiration, ATP production, and spare respiratory capacity, respectively. Data are presented as mean ± SEM (n = 3). \*\**P* < 0.01, \*\*\**P* < 0.001, \*\*\*\**P* < 0.0001; ns, not significant.

Collectively, these findings indicate that FB preferentially attenuates LPS-induced glycolytic activation and partially improves basal mitochondrial respiration, while exerting limited effects on other mitochondrial respiratory parameters.

### 3.4. FB promotes HIF-1α ubiquitination and proteasomal degradation and suppresses HIF-1α-dependent glycolytic and inflammatory responses

To determine whether FB regulated HIF-1α at the transcriptional level, *Hif1a* mRNA expression was measured in L4–L6 spinal cord tissues and BV2 microglia. CCI increased *Hif1a* mRNA expression in spinal cord tissues compared with the Sham group, whereas FB treatment did not significantly change this increase (Fig. 4A). Similarly, LPS increased *Hif1a* mRNA expression in BV2 microglia, but FB treatment did not significantly affect the LPS-induced increase (Fig. 4B). These findings suggest that FB mainly regulated HIF-1α at the post-transcriptional level.

**Fig. 4.**
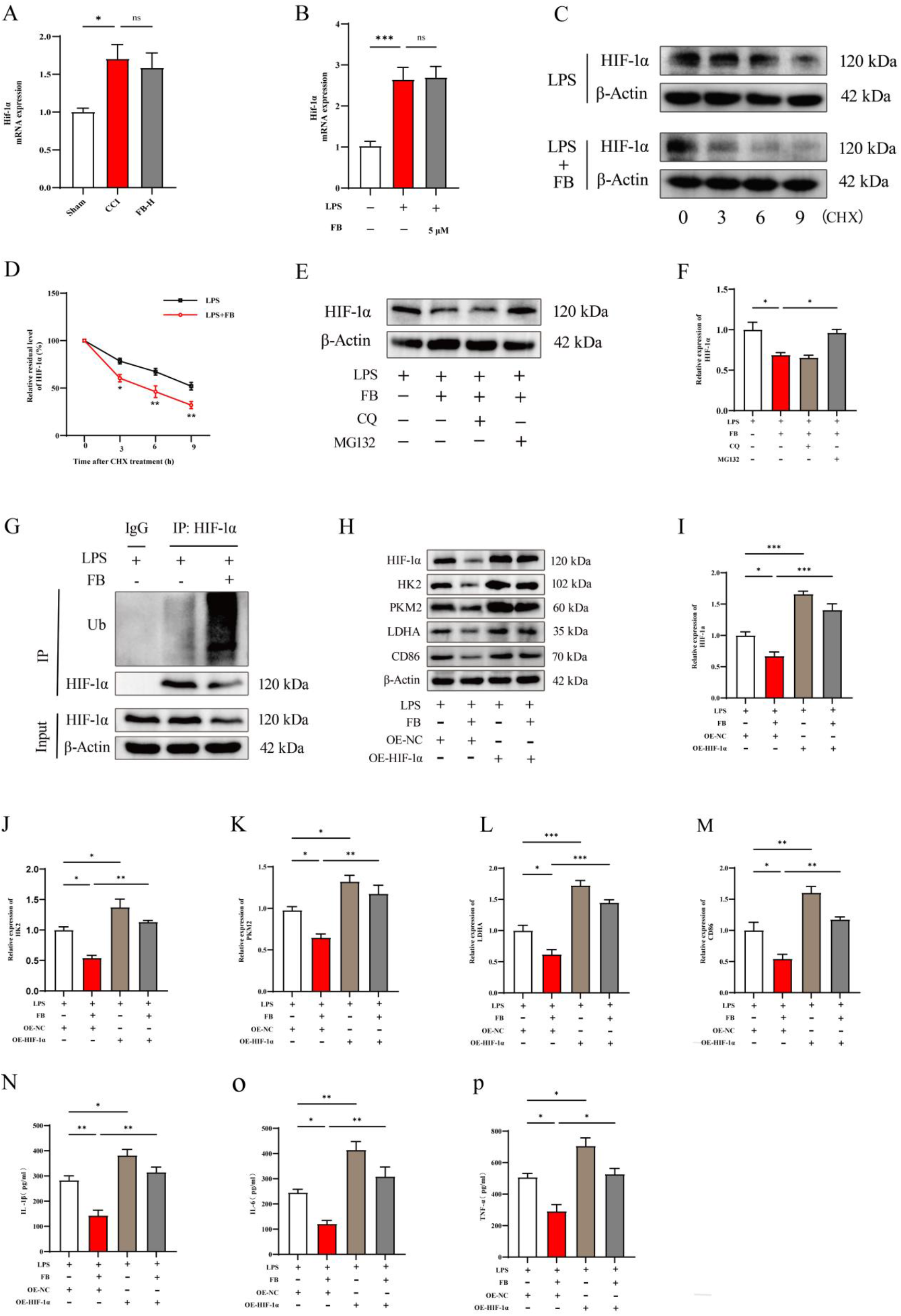
FB promotes HIF-1α degradation and suppresses glycolytic and inflammatory responses. (A) *Hif1a* mRNA expression in L4–L6 spinal cord tissues from the Sham, CCI, and FB-H groups. (B) *Hif1a* mRNA expression in BV2 microglia exposed to LPS and treated with 5 μM FB. (C) Representative Western blot images from the CHX chase assay showing HIF-1α protein levels in LPS-stimulated BV2 microglia treated with or without FB at the indicated times after CHX administration. (D) Quantification of HIF-1α protein levels in the CHX chase assay. (E) Representative Western blot images of HIF-1α after treatment with FB in the presence or absence of the lysosomal inhibitor CQ or the proteasome inhibitor MG132. (F) Quantification of HIF-1α protein levels shown in E. (G) HIF-1α ubiquitination analyzed by immunoprecipitation with an anti-HIF-1α antibody followed by immunoblotting with an anti-ubiquitin antibody. (H) Representative Western blot images of HIF-1α, HK2, PKM2, LDHA, and CD86 in LPS-stimulated BV2 cells treated with FB and transfected with a negative-control vector or HIF-1α-overexpression vector. (I–M) Quantification of HIF-1α, HK2, PKM2, LDHA, and CD86 protein levels, respectively, normalized to β-actin. (N–P) IL-1β, IL-6, and TNF-α levels, respectively, in culture supernatants measured by ELISA. Data are presented as mean ± SEM (n = 3). \**P* < 0.05, \*\**P* < 0.01, and \*\*\**P* < 0.001.

CHX chase analysis showed that FB accelerated HIF-1α degradation in LPS-stimulated BV2 microglia (Fig. 4C,D). MG132, but not chloroquine, prevented the FB-induced reduction in HIF-1α protein levels (Fig. 4E,F). Consistent with these findings, immunoprecipitation followed by immunoblotting showed that FB increased HIF-1α ubiquitination (Fig. 4G).

To assess the contribution of HIF-1α to the effects of FB, we overexpressed HIF-1α in LPS-stimulated BV2 cells. HIF-1α overexpression partly reversed the FB-induced reductions in HIF-1α, HK2, PKM2, LDHA, and CD86 expression (Fig. 4H–M). It also weakened the inhibitory effects of FB on IL-1β, IL-6, and TNF-α production (Fig. 4N– P). Overall, these findings support a role for FB-induced HIF-1α degradation in the suppression of glycolytic and inflammatory responses in microglia.

### 3.5. USP7 is identified as a candidate deubiquitinating enzyme involved in HIF-1α regulation in inflammatory microglia

To identify candidate regulators of HIF-1α ubiquitination, we performed HIF-1α immunoprecipitation followed by mass spectrometry (IP-MS) in LPS-stimulated BV2 microglia. Among the 57 proteins annotated as E3 ubiquitin ligases or deubiquitinating enzymes in the IP-MS dataset, USP7 was the only protein shared with the 15 HIF-1α regulators retrieved from UbiBrowser 2.0 (Fig. 5A; Supplementary Tables 2 and 3). USP7 was identified by 26 peptides, including 20 unique peptides. Therefore, we selected USP7 for further validation.

**Fig. 5.**
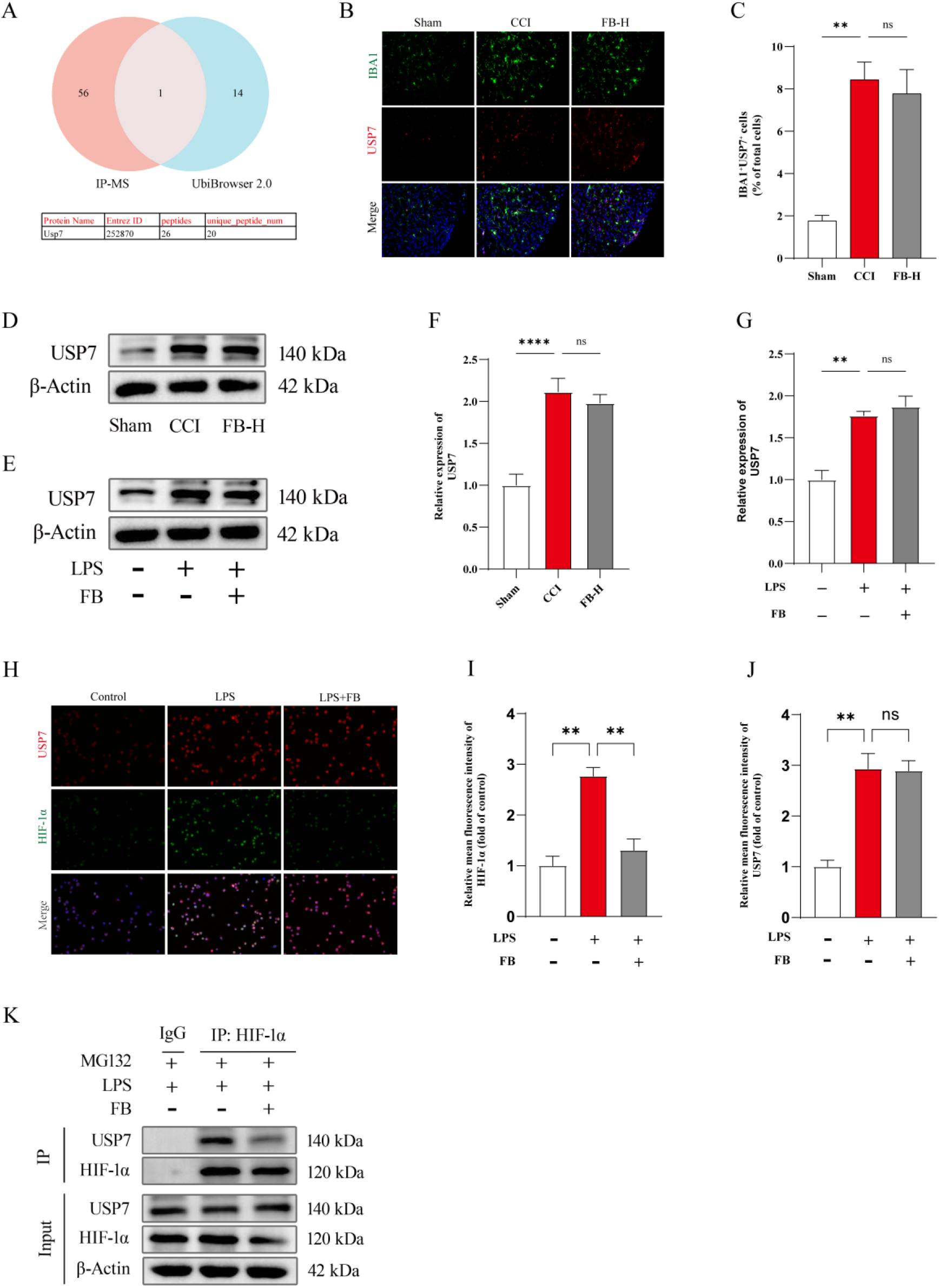
Identification and validation of USP7 as a candidate regulator of HIF-1α in inflammatory microglia. (A) Overlap between E3 ubiquitin ligases and deubiquitinating enzymes identified by HIF-1α immunoprecipitation followed by mass spectrometry (IP-MS) and HIF-1α regulators retrieved from UbiBrowser 2.0. The inset table lists the peptide identification data for USP7. (B) Representative immunofluorescence images of IBA1 and USP7 in the L4–L6 spinal dorsal horn of Sham, CCI, and FB-H mice. Scale bar, 100 μm. (C) Quantification of IBA1⁺/USP7⁺ cells. (D,E) Representative Western blot images of USP7 in L4–L6 spinal cord tissues and BV2 microglia, respectively. (F,G) Quantification of USP7 protein levels in spinal cord tissues and BV2 cells, respectively, normalized to β-actin. (H) Representative immunofluorescence images of USP7 and HIF-1α in BV2 cells. Scale bar, 50 μm. (I,J) Quantification of HIF-1α and USP7 fluorescence intensity, respectively. (K) Co-IP analysis of the association between USP7 and HIF-1α in LPS-stimulated BV2 cells treated with MG132, with or without FB. Data are presented as mean ± SEM (n = 3). \*\**P* < 0.01 and \*\*\*\**P* < 0.0001; ns, not significant.

In the spinal dorsal horn, CCI increased the proportion of IBA1⁺/USP7⁺ cells, whereas high-dose FB did not significantly reduce this increase (Fig. 5B,C). Consistent with this finding, CCI increased USP7 protein expression in L4–L6 spinal cord tissues. However, USP7 levels did not differ significantly between the CCI and FB-H groups (Fig. 5D,F).

LPS also increased USP7 protein expression in BV2 microglia, whereas FB treatment did not significantly change total USP7 abundance (Fig. 5E,G). Immunofluorescence analysis further showed that FB reduced HIF-1α fluorescence intensity without affecting USP7 fluorescence intensity in LPS-stimulated BV2 microglia (Fig. 5H–J). These findings suggest that the inhibitory effect of FB on HIF-1 α was not caused by a reduction in total USP7 abundance.

Finally, Co-IP analysis showed an association between USP7 and HIF-1α in LPS-stimulated BV2 cells. FB treatment weakened this association (Fig. 5K). Overall, USP7 was identified as a candidate deubiquitinase involved in HIF-1 α regulation. The findings also suggest that FB may promote HIF-1α degradation by weakening the association between USP7 and HIF-1α.

### 3.6. FB engages USP7 and modulates HIF-1α ubiquitination and inflammatory activation in BV2 microglia

To examine the potential interaction between FB and USP7, we performed molecular docking using the USP7 structure with PDB ID 5VSB. The docking analysis predicted that FB could occupy a binding pocket in USP7, with a docking score of −7.827 kcal/mol (Fig. 6A). The predicted binding mode involved GLN293, ASP295, ARG408, PHE409, TYR411, ASN418, and TYR514, with possible hydrogen-bond and aromatic interactions. CETSA showed that FB increased the thermal stability of USP7. Compared with vehicle treatment, FB increased the levels of residual USP7 protein at 50–60 °C (Fig. 6B,C). These findings support a potential interaction between FB and USP7 in BV2 microglia.

**Fig. 6.**
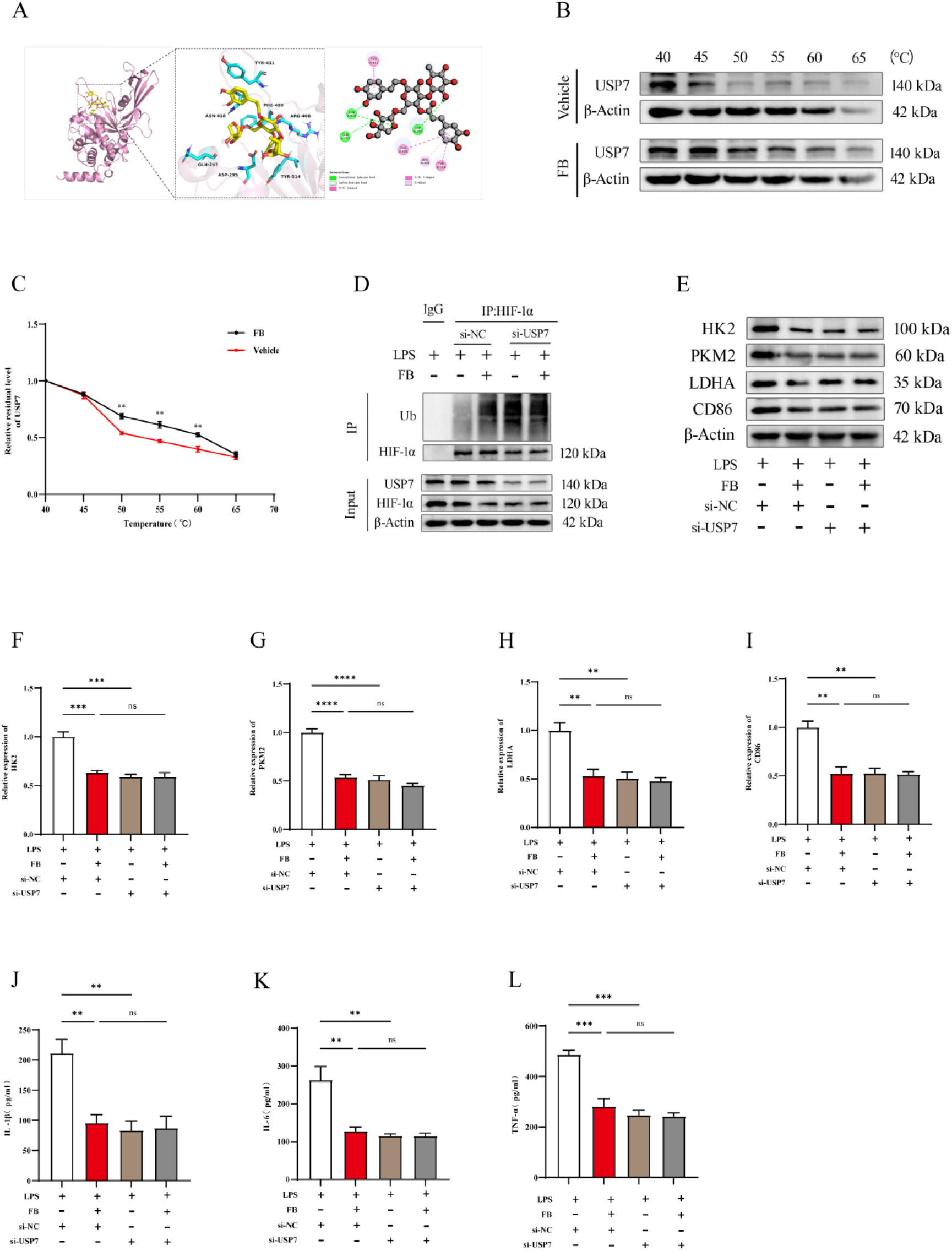
FB engages USP7 and modulates HIF-1α ubiquitination and inflammatory activation in BV2 microglia. Predicted binding mode and two-dimensional interaction map of FB in the USP7 binding pocket. (B) Representative Western blot images from the CETSA. (C) Quantification of residual USP7 levels in the CETSA. (D) HIF-1α immunoprecipitation followed by immunoblotting for ubiquitin in LPS-stimulated BV2 microglia treated with FB, si-USP7, or both. USP7, HIF-1α, and β-actin levels in the input samples are shown. (E) Representative Western blot images of HK2, PKM2, LDHA, and CD86. (F–I) Quantification of HK2, PKM2, LDHA, and CD86 protein levels, respectively, normalized to β-actin. (J–L) IL-1β, IL-6, and TNF-α concentrations in culture supernatants, respectively, measured by ELISA. Data are presented as mean ± SEM (n = 3). \*\**P* < 0.01, \*\*\**P* < 0.001, and \*\*\*\**P* < 0.0001; ns, not significant.

We next evaluated the effects of USP7 knockdown. Among the three USP7-targeting siRNAs, si-USP7-3 showed the highest knockdown efficiency and was selected for subsequent experiments (Supplementary Fig. 1A,B). Immunoprecipitation of HIF-1α followed by immunoblotting for ubiquitin showed that both FB treatment and USP7 knockdown increased HIF-1α ubiquitination. The combined treatment did not produce a further increase, suggesting that FB and USP7 knockdown had overlapping effects on HIF-1α ubiquitination (Fig. 6D). FB treatment and USP7 knockdown each reduced the protein levels of HK2, PKM2, LDHA, and CD86 in LPS-stimulated BV2 microglia (Fig. 6E–I). The combined treatment did not further reduce these protein levels. Similarly, FB treatment and USP7 knockdown reduced the secretion of IL-1β, IL-6, and TNF-α. The combined treatment did not produce additional effects (Fig. 6J–L).

Overall, USP7 knockdown reproduced the effects of FB on HIF-1α ubiquitination, glycolysis-related protein expression, and inflammatory cytokine production. The absence of an additional effect after combined treatment is consistent with the involvement of USP7 in the cellular responses regulated by FB.

## 4. Discussion

In the present study, FB alleviated mechanical allodynia and thermal hyperalgesia in CCI mice. This behavioral improvement was accompanied by reduced expression of HIF-1α, HK2, PKM2, LDHA, and CD86 in the L4–L6 spinal cord. FB also decreased the proportion of IBA1+/CD86+ microglia and reduced spinal IL-1β, IL-6, and TNF-α levels. These coordinated changes suggest that the analgesic effect of FB is associated with reduced spinal neuroinflammation, proinflammatory microglial responses, and glycolytic alterations. Spinal microglia contribute to pain hypersensitivity through inflammatory mediators that enhance nociceptive transmission [24,25]. In addition, metabolic reprogramming is closely linked to microglial phenotypic changes and inflammatory responses during neuropathic pain [26]. The present findings are therefore consistent with a role for microglial metabolic regulation in the analgesic action of FB.

To examine this possibility in microglia, we evaluated FB in LPS-stimulated BV2 cells. LPS induces coordinated inflammatory and metabolic responses in murine and human microglia, including enhanced glycolytic activity and mitochondrial dysfunction [27–29]. Consistent with these observations, LPS increased HIF-1α, HK2, PKM2, LDHA, and CD86 expression in BV2 microglia, together with the secretion of IL-1β, IL-6, and TNF-α. FB reduced each of these responses. The concurrent reductions in glycolytic enzymes, CD86, and cytokine secretion indicate that FB suppresses glycolytic alterations and proinflammatory activation in microglia. These cellular results were consistent with the metabolic and inflammatory changes observed in the spinal cord of CCI mice.

After establishing the effects of FB on the inflammatory phenotype, we assessed glycolytic and mitochondrial functions using extracellular flux analysis. LPS increased glycolysis, glycolytic capacity, and glycolytic reserve, whereas FB significantly attenuated these changes. FB also partially restored basal mitochondrial respiration, while maximal respiration, ATP production, and spare respiratory capacity remained unchanged relative to the LPS group. This pattern suggests that FB preferentially suppresses excessive glycolytic activity while partially improving basal mitochondrial respiration. A similar increase in ECAR accompanied by reduced OCR has been reported in LPS treated primary microglia [30]. Moreover, enhanced microglial glycolysis has been linked to neuroinflammation and pain hypersensitivity in CCI and diabetic neuropathic pain models [31–34]. Thus, the reduction in glycolytic flux induced by FB may contribute to the attenuation of proinflammatory microglial responses and spinal pain sensitization.

We next examined whether FB regulated HIF-1α expression and protein stability. FB did not significantly reduce the increases in *Hif1a* mRNA induced by CCI or LPS. However, FB accelerated HIF-1α protein degradation in the cycloheximide chase assay and increased HIF-1α ubiquitination. MG132, but not chloroquine, prevented the FB-induced reduction in HIF-1α protein. These findings support the involvement of ubiquitin–proteasome-dependent HIF-1α degradation rather than lysosomal degradation. HIF-1α is closely associated with glycolytic programming and inflammatory signaling in microglia [35–37]. Accordingly, reduced HIF-1α stability provides a mechanistic basis for the decreases in HK2, PKM2, and LDHA expression observed after FB treatment.

The functional contribution of HIF-1α was further assessed by overexpression. HIF-1α overexpression partially reversed the FB-induced reductions in HK2, PKM2, LDHA, and CD86 expression and attenuated the reductions in IL-1β, IL-6, and TNF-α secretion. These data support a role for HIF-1α protein stability in the glycolytic and proinflammatory responses regulated by FB. Collectively, the animal and cellular findings support a model in which FB promotes HIF-1α degradation through the ubiquitin–proteasome pathway, thereby limiting HIF-1α-associated glycolytic reprogramming and proinflammatory activation in microglia. This interpretation is consistent with evidence that microglial metabolic regulation influences neuroinflammation and neuropathic pain outcomes [6,38].

Because FB increased HIF-1α ubiquitination and promoted its proteasomal degradation without reducing *Hif1a* mRNA expression, we next investigated the upstream posttranslational mechanisms that may regulate HIF-1α protein stability. Ubiquitination and deubiquitination constitute a reversible regulatory system that controls protein stability and signalling output. In neuroinflammatory conditions, ubiquitin modifying enzymes regulate the turnover and activity of inflammatory signalling proteins, thereby shaping microglial functional states [39,40]. Deubiquitinating enzymes can also counteract ubiquitin dependent proteasomal turnover of HIF proteins and influence the magnitude and duration of HIF signalling [41].

USP7 is a multifunctional deubiquitinase that regulates the stability of diverse substrate proteins and has emerged as a potential pharmacological target [42]. Molecular docking predicted that FB could occupy a binding pocket in USP7, and CETSA showed increased thermal stability of USP7 in FB-treated BV2 microglia. Together, these findings support USP7 as a potential intracellular target of FB. USP7 was the only candidate shared by the HIF-1α immunoprecipitation–mass spectrometry dataset and UbiBrowser analysis. Although FB did not alter total USP7 protein expression, it weakened the association between USP7 and HIF-1α. In addition, FB treatment and USP7 knockdown produced comparable increases in HIF-1α ubiquitination and reductions in HK2, PKM2, LDHA, CD86, and proinflammatory cytokine secretion. Combined treatment produced no further effect. These findings are consistent with the involvement of USP7 in the actions of FB. They further suggest that FB may interfere with USP7-dependent maintenance of HIF-1α stability, thereby facilitating HIF-1α ubiquitination and ubiquitin–proteasome-dependent degradation. This regulatory effect may subsequently limit HIF-1α-associated glycolytic reprogramming and proinflammatory activation in microglia.

Several limitations should be considered. First, only the CCI model and LPS-stimulated BV2 cells were used. These models do not fully reflect the heterogeneity of neuropathic pain or the responses of primary spinal microglia after peripheral nerve injury. Second, molecular docking and CETSA support a potential interaction between FB and USP7, but do not establish direct binding. The binding site of FB on USP7, the critical residues involved, and the effect of FB on USP7 catalytic activity require further study. Third, the requirement for spinal microglial USP7 in FB-mediated analgesia was not examined in vivo. Microglia-specific manipulation of USP7 in CCI mice would help define the causal contribution of this pathway. Finally, ECAR and OCR were evaluated only in BV2 cells, and the pharmacokinetic profile, spinal exposure, and long-term safety of FB remain to be determined.

Despite these limitations, the present findings support the involvement of the USP7–HIF-1α axis in the regulation of microglial glycolytic and proinflammatory responses by FB. These data provide a mechanistic basis for further evaluating USP7 as a potential target for neuropathic pain.

## 5. Conclusion

In this study, FB attenuated mechanical allodynia and thermal hyperalgesia in CCI mice and reduced spinal neuroinflammation, accompanied by decreased HIF-1α-associated glycolytic markers and proinflammatory microglial activation. Complementary studies in LPS-stimulated BV2 microglia showed that FB suppressed glycolytic and proinflammatory responses and promoted HIF-1 α ubiquitination and proteasome-dependent degradation. FB may interact with USP7 and weaken the USP7-HIF-1α association, thereby facilitating HIF-1 α degradation. Collectively, these findings suggest that FB may alleviate neuropathic pain by limiting HIF-1 α -associated microglial glycolytic reprogramming through modulation of the USP7/HIF-1α axis.

## Funding

This work was supported by the Applied Basic Research Program (Grant No. 2022JH2/101300019).

## Data availability

The data supporting the findings of this study are available from the corresponding author upon reasonable request.

### CRediT authorship contribution statement

Zhen Dai: Conceptualization, Methodology, Data curation, Writing – original draft, Writing – review & editing.

Hao Wang: Formal analysis, Data curation, Writing – review & editing. Yanfeng Yang: Supervision, Project administration, Writing – review & editing.

### Declaration of competing interest

The authors declare no competing interests.

## Notes

### Competing Interest Statement

The authors have declared no competing interest.

