## Supplementary_Fig_1 for "Forsythoside B attenuates neuropathic pain by suppressing USP7/HIF-1α-mediated microglial glycolytic reprogramming"

**Supplementary Fig. 1. Selection of an effective siRNA targeting USP7 in BV2 microglia.**

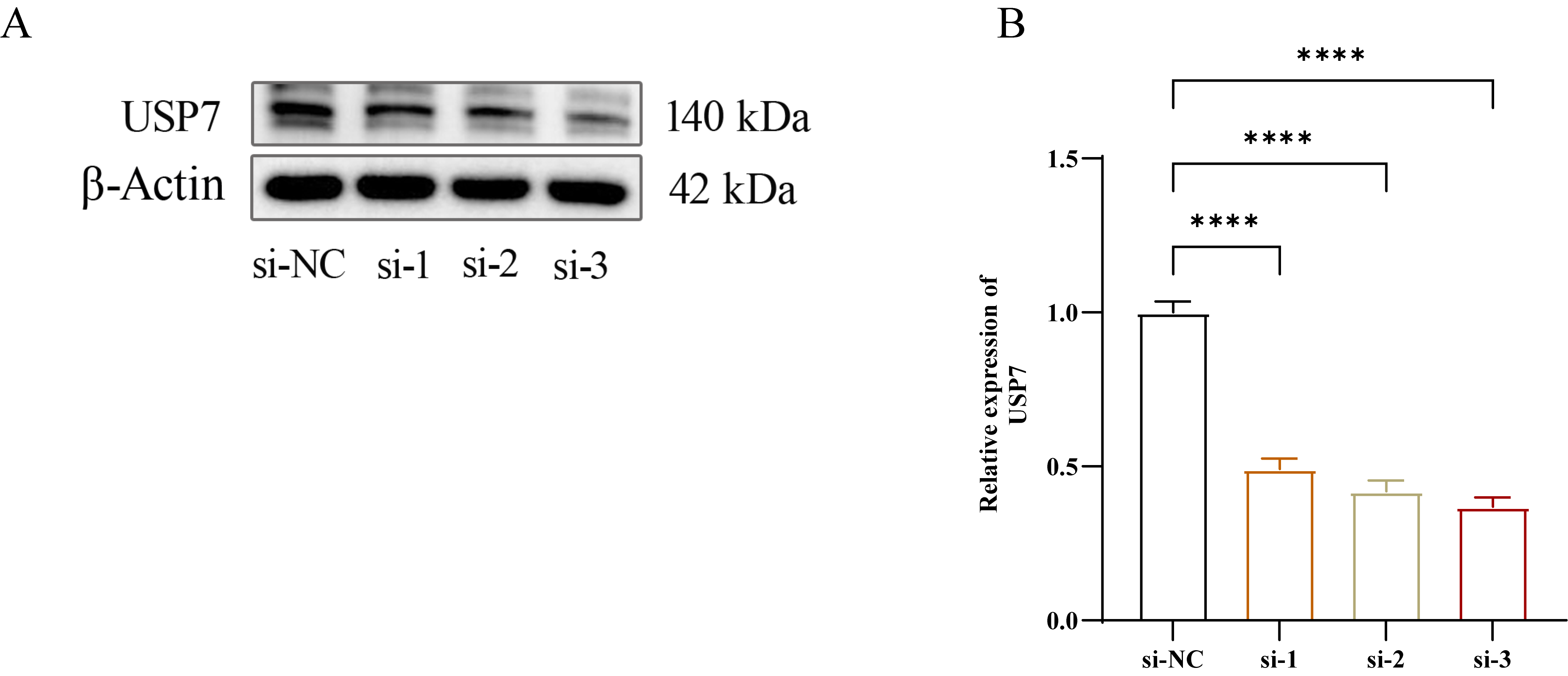

(A) Representative western blot images of USP7 in BV2 microglia transfected with negative control siRNA or three siRNAs targeting USP7. (B) Quantification of USP7 protein expression normalized to β-actin. Data are presented as the mean ± SEM (n = 3). *****P* < 0.0001.
